# Squidpy: a scalable framework for spatial single cell analysis

**DOI:** 10.1101/2021.02.19.431994

**Authors:** Giovanni Palla, Hannah Spitzer, Michal Klein, David Fischer, Anna Christina Schaar, Louis Benedikt Kuemmerle, Sergei Rybakov, Ignacio L. Ibarra, Olle Holmberg, Isaac Virshup, Mohammad Lotfollahi, Sabrina Richter, Fabian J. Theis

**Affiliations:** Institute of Computational Biology, Helmholtz Center Munich, Germany; TUM School of Life Sciences Weihenstephan, Technical University of Munich, Germany; Department of Mathematics, Technical University of Munich, Germany; Institute for Tissue Engineering and Regenerative Medicine (iTERM), Helmholtz Center Munich, Germany; Department of Anatomy and Physiology, University of Melbourne, Australia

## Abstract

Spatial omics data are advancing the study of tissue organization and cellular communication at an unprecedented scale. Here, we present Squidpy, a Python framework that brings together tools from omics and image analysis to enable scalable description of spatial molecular data, such as transcriptome or multivariate proteins. Squidpy provides both infrastructure and numerous analysis methods that allow to efficiently store, manipulate and interactively visualize spatial omics data.

## Main

Dissociation-based single cell technologies have enabled the deep characterization of cellular states and the creation of cell atlases of many organs and species^1^. However, how cellular diversity constitutes tissue organization and function is still an open question. Spatially-resolved molecular technologies aim at bridging this gap by enabling the investigation of tissues in situ at cellular and subcellular resolution^2-4^. In contrast to the current state of the art dissociation-based protocols, spatial molecular technologies acquire data in greatly diverse forms, in terms of resolution (few cells per observation to subcellular resolution), multiplexing (dozens of features to genome-wide expression profiles), modality (transcriptomics, proteomics and metabolomics) and often times with an associated high-content image of the captured tissue^2-4^. Such diversity in resulting data and corresponding formats currently represents an organisational hurdle that has hampered urgently needed development of interoperable and broad analysis methods. The underlying computational challenge requires solutions both in terms of efficient data representation as well as comprehensive analysis and visualization methods.

Hence, existing analysis frameworks for spatial data focus either on pre-processing^5-8^ or on one particular aspect of spatial data analysis^9-13^. Due to the lack of a unified data representation and modular API, users so far cannot perform comprehensive analyses leveraging the full spatial modality, e.g. combining stlearn’s^12^ integrative analysis of tissue images together with Giotto‘;s powerful spatial statistics^11^. A comprehensive framework that enables community-driven scalable analysis of both spatial neighborhood graph and image, along with an interactive visualization module, is missing (Supplementary Table 1).

For this purpose we developed “Spatial Quantification of Molecular Data in Python” (Squidpy), a python-based framework for the analysis of spatially-resolved omics data (Fig. 1). Squidpy aims to bring the diversity of spatial data in a common data representation and provide a common set of analysis and interactive visualization tools. Such infrastructure is useful in a variety of analysis settings, for different data types, and it explicitly leverages the additional information that spatial data provides: the spatial coordinates and, when available, the tissue image. Squidpy is built on top of Scanpy and Anndata^14^, and it relies on several scientific computing libraries in Python, such as Scikit-image^15^ and Napari^16^. Its modularity makes it suitable to be interfaced with a variety of additional tools in the python data science and machine learning ecosystem, as well as several single-cell data analysis packages. It allows to quickly explore spatial datasets and lays the foundations for both spatial omics data analysis as well as novel methods development.

**Figure 1:**
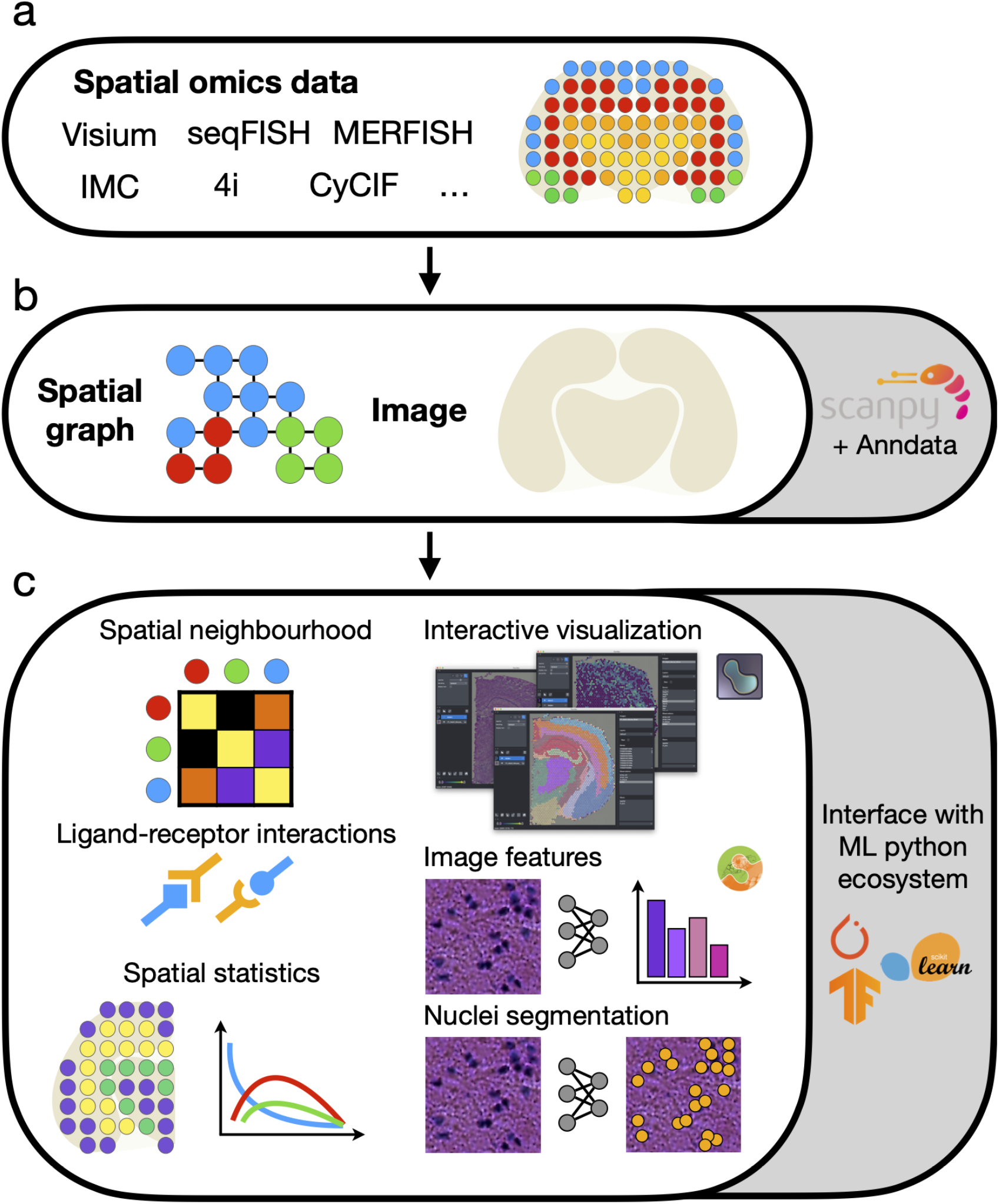
Squidpy is a software framework for the analysis of spatial omics data. (a) Squidpy supports inputs from diverse spatial molecular technologies with spot-based, single-cell, or subcellular spatial resolution. (b) Building upon the single-cell analysis software Scanpy^14^ and the Anndata format, Squidpy provides efficient data representations of these inputs, storing spatial distances between observations in a spatial graph and providing an efficient image representation for high resolution tissue images that might be obtained together with the molecular data. (c) Using these representations, several analysis functions are defined to quantitatively describe tissue organization at cellular (spatial neighborhood) and gene level (spatial statistics, spatially-variable genes and ligand-receptor interactions), to combine microscopy image information (image features and nuclei segmentation) with omics information and to interactively visualize high-resolution images.

## Results

### Squidpy provides technology-agnostic data representations for spatial graphs and images

Squidpy introduces two main data representations to manage and store spatial omics data in a technology-agnostic way: a neighborhood graph from spatial coordinates, and large source images acquired in spatial omics data (Fig. 1b). Spatial graphs encode spatial proximity, and are, depending on data resolution, flexible in order to support the variety of neighborhood metrics that spatial data types and users may require. For instance, in Spatial Transcriptomics (ST^17^, Visium^18^, DBit-seq^19^), a node is a spot and a neighborhood set can be defined by a fixed number of adjacent spots whereas in imaging-based molecular data (seqFISH^20^, MERFISH^21^, Imaging Mass Cytometry^22,23^, CyCif^24^, 4i^25^, Spatial Metabolomics^26^, see Fig. 1a), a node can be defined as a cell (or pixel), and a neighborhood set can also be chosen based on a fixed radius (expressed in spatial units) from the centroid of each observation. Alternatively, other dissimilarity measures, such as euclidean distance, can be utilized to build the neighbor graph. Such data representation is suitable for many analysis tools that aim at quantifying spatial organization of the tissue. In Squidpy, we provide several tools to compute statistics at cell and gene level, such as a neighborhood enrichment score on the spatial graph, a ligand-receptor interaction analysis tool, and the Moran’s I spatial autocorrelation score for spatially variable genes identification (Fig. 1c).

The high resolution microscopy image additionally captured by spatial omics technologies represents a rich source of morphological information that can provide key biological insights into tissue structure and cellular variation. Squidpy introduces a new data object, the Image Container, that efficiently stores the image with an on-disk/in-memory switch based on xArray and Dask^27,28^. The Image Container provides image analysis tools, such as performing image preprocessing, segmentation, and feature extraction, as well as interfacing with modern deep learning frameworks for more advanced analysis^15^ (Fig. 1c right). It provides seamless integration with Napari^16^, thus enabling interactive visualization of analysis results stored in an Anndata object alongside the high resolution image directly from a Jupyter notebook. It also enables interactive manual cropping of tissue areas and automatic annotation of observations in Anndata. Since Napari is an image viewer in Python, all the above-mentioned functionalities can be also interactively executed without additional requirements.

### Squidpy enables calculation of spatial cellular statistics using spatial graphs

A key question in the analysis of spatial molecular data is the description and quantification of spatial patterns and cellular neighborhoods across the tissue. Squidpy provides several tools that leverage the spatial graph to address such questions. For instance, a neighborhood enrichment analysis score that quantifies cluster proximity with a permutation based test (see online methods) is available. When applied to a recently published seqFISH data of mouse gastrulation^29^, we found several clusters to be co-enriched in their cellular neighbors (Fig. 2a,b), recapitulating the main results of the original authors. Furthermore, our implementation is scalable and ∼10 fold faster than a similar implementation in Giotto^11^ (Supplementary Fig. 1a), enabling analysis of large-scale spatial omics datasets. Squidpy also computes a co-occurrence score for clusters across spatial coordinates, which we applied to a 4i dataset of Hela cells^25^. We considered ∼270,000 pixels as subcellular resolution observations, and evaluated their cluster co-occurrence at increasing distances (Fig. 2 c,d). As expected, the subcellular measurements annotated in the Nucleus compartment co-occur together with the Nucleus and the Nuclear envelope, at short distances. Squidpy provides additional tools to investigate features of spatial-molecular data, such as a fast and broader implementation of CellPhoneDB^30^ for spatial ligand-receptor interaction analysis, leveraging the larger Omnipath database^31^, and the Moran’s I spatial autocorrelation statistic for detection of spatially variable genes^32^ (Fig 2 e,f). These statistics yield interpretable results across diverse experimental techniques, as we demonstrate on an Imaging Mass Cytometry dataset^33^, where we showcase additional methods like the Ripley’s K function, average clustering, and degree and closeness centrality (see Supplementary Fig. 3).

**Figure 2:**
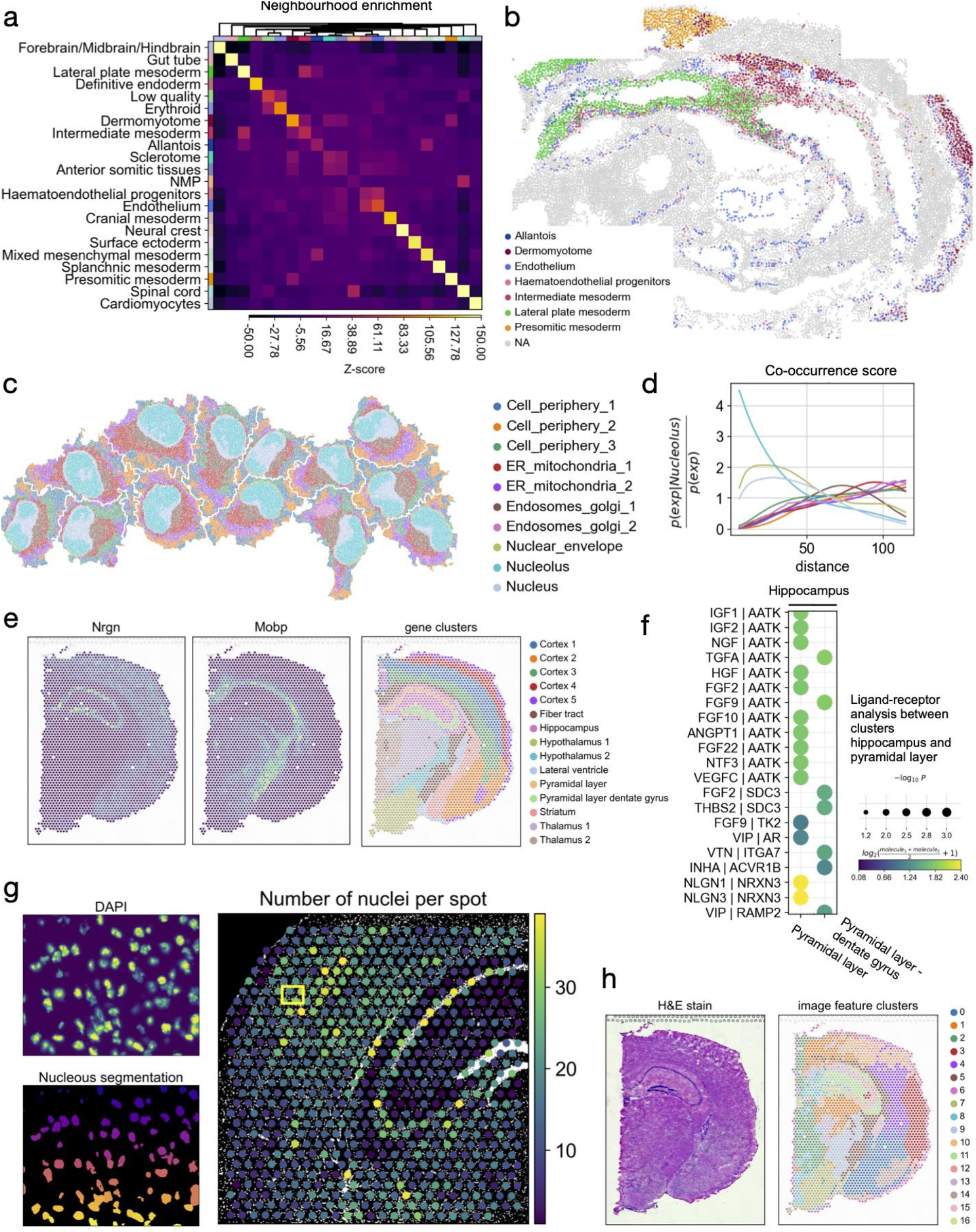
Analysis of spatial omics datasets across diverse experimental techniques using Squidpy. (a) Neighborhood enrichment analysis between cell clusters in spatial coordinates. The “Lateral plate mesoderm” cluster is co-enriched with the “Alllantois” and “Intermediate mesoderm” cluster. Also, the “Endothelium” cluster is enriched with the “Haematoendothelial progenitors”. Both of these results were also reported by the original authors^29^. (b) Visualization of selected clusters of the seqFISH mouse gastrulation dataset. (c) Visualization of subcellular molecular profiles in Hela Cells, plotted in spatial coordinates (approx 270000 observations/pixels). (d) Cluster co-occurrence score at increasing distance threshold across the tissue. The cluster “Nucleolus” is found to be co-enriched at short distances with the “Nucleus” and the “Nuclear envelope” clusters. (e) Expression of Nrgn, Mobp, and clustering result from gene expression space plotted on spatial coordinates. Nrgn and Mobp are spatially variable genes defined with Moran’s I global spatial autocorrelation score. The selected genes are spatially distributed and they are shared across different clusters. (f) Ligand-receptor interactions from the cluster “Hippocampus” to clusters “Pyramidal Layer” and “Pyramidal layer dentate gyrus”. Shown are a subset of significant ligand-receptor pairs queried using Omnipath database. (g) Segmentation features derived from fluorescence image of Visium mouse brain dataset. Top left: DAPI stain. Bottom left: nuclei segmentation using DAPI stain. Right: number of nuclei in each Visium spot derived from the nuclei segmentation count. The yellow square shows the location of the inset. (h) H&E stain and clustering of summary image features (channel intensity mean, standard deviation, and 0.1, 0.5, 0.9th quantiles) derived from the H&E stain at each spot location (for quantitative comparison see Supplementary Fig. 2e).

### Squidpy allows analysis of images in spatial omics analysis workflows

Squidpy’s Image Container object provides a general mapping between pixel coordinates and molecular profile, enabling analysts to relate image-level observations to omics measurements.

Following standard image-base profiling techniques^34^, Squidpy implements a pipeline based on Scikit-image^15^ for preprocessing and segmenting images, extracting morphological, texture, and deep learning-powered features (Supplementary Fig. 2a). To enable efficient processing of very large images, this pipeline utilises lazy loading, image tiling and multi-processing (Supplementary Fig. 1b). Features can be extracted from a raw tissue image crop, or Squidpy’s nuclei-segmentation module can be used to extract nuclei counts and nuclei sizes (Supplementary Fig. 2b). For instance, we can leverage segmented nuclei to inform cell-type deconvolution methods such as Tangram^35^ or Cell2Location^36^ (Supplementary Fig. 4).

As an example of segmentation-based features, we calculated a nuclei segmentation using the DAPI stain of a fluorescence mouse brain section and showed the estimated number of nuclei per spot on the hippocampus (Fig. 2g). The cell-dense pyramidal layer can be easily distinguished with this view of the data, showcasing the richness and interpretability of information that can be extracted from tissue images when brought in a spot-based format.

Squidpy’s feature extraction pipeline enables direct comparison and joint analysis of image data and omics data. For instance, using a Visium mouse brain dataset, we compared gene clusters with a clustering of summary features (mean, standard deviation, 0.1, 0.5, and 0.9th quantiles) of the accompanying H&E stained tissue image (Fig. 2e,h). Several image feature clusters show similarities with the gene-based clusters, especially in the hippocampus (77% overlap with image feature cluster 10), and the hypothalamus (54% overlap with image feature cluster 10), but provide a different view of the data in the cortex (no overlap >33% with any image feature clusters) (Supplementary Fig. 2e).

## Conclusion

In summary, Squidpy enables the analysis of spatial molecular data by leveraging two data representations: the spatial graph and the tissue image. It interfaces with Scanpy and the Python data science ecosystem, providing a scalable and extendable framework for novel methods development in the field of biological spatial molecular data. We are convinced that Squidpy could contribute to building a bridge between the molecular omics community and the image analysis and computer vision community to develop the next generation of computational methods for spatial omics technologies.

## Code and data availability

Squidpy is a pip installable python package and available at the following github repository: https://github.com/theislab/squidpy, with documentation at: https://squidpy.readthedocs.io/en/latest/. All the results of this analysis can be found at the following github repository: https://github.com/theislab/squidpy_reproducibility. The pre-processed datasets have been deposited at https://doi.org/10.6084/m9.figshare.c.5273297.v1 and they are all conveniently accessible in Python via the *squidpy*.*dataset* module. The datasets used in this article are the following: Imaging Mass Cytometry^33^, seqFISH^29^, 4i^25^, and several Visium^18^ datasets available from the website: https://support.10xgenomics.com/spatial-gene-expression/datasets.

## Acknowledgments

We would like to acknowledge Luke Zappia, Malte Luecken and all the members of Theis lab for helpful discussion. We would like to acknowledge Gemma Fornons for the Squidpy logo, Scanpy developers Philipp Angerer and Fidel Ramirez for useful discussion and code revision. We would like to thank authors of original publications and 10x Genomics for making spatial omics datasets publicly available.

Sa.R., G.P. are supported by the Helmholtz Association under the joint research school “Munich School for Data Science” - MUDS.

A.C.S. has been funded by the German Federal Ministry of Education and Research (BMBF) under Grant No. 01IS18036B.

F.J.T. acknowledges support by the BMBF (grant# 01IS18036B, grant# 01IS18053A and grant# 031L0210A), the European Union’s Horizon 2020 research and innovation programme under grant agreement No 874656, the Chan Zuckerberg Initiative DAF (advised fund of Silicon Valley Community Foundation, grant # 2019-207271), the Bavarian Ministry of Science and the Arts in the framework of the Bavarian Research Association “ForInter” (Interaction of human brain cells) and by the Helmholtz Association’s Initiative and Networking Fund through Helmholtz AI [grant number: ZT-I-PF-5-01] and sparse2big [grant number ZT-I-007].

F.J.T. reports receiving consulting fees from Roche Diagnostics GmbH and Cellarity Inc., and ownership interest in Cellarity, Inc. and Dermagnostix

## Author contributions

G.P., H.S., M.K., D.F., F.J.T. designed the study. G.P., H.S., M.K., D.F., A.C.S, L.B.K., Se.R., I.L.I., O.H., I.V., M.L., Sa.R. wrote the code. G.P., H.S., M.K. performed the analysis. F.J.T. supervised the work. All authors read and corrected the final manuscript.

## Supplements

**Supplementary Figure 1.**
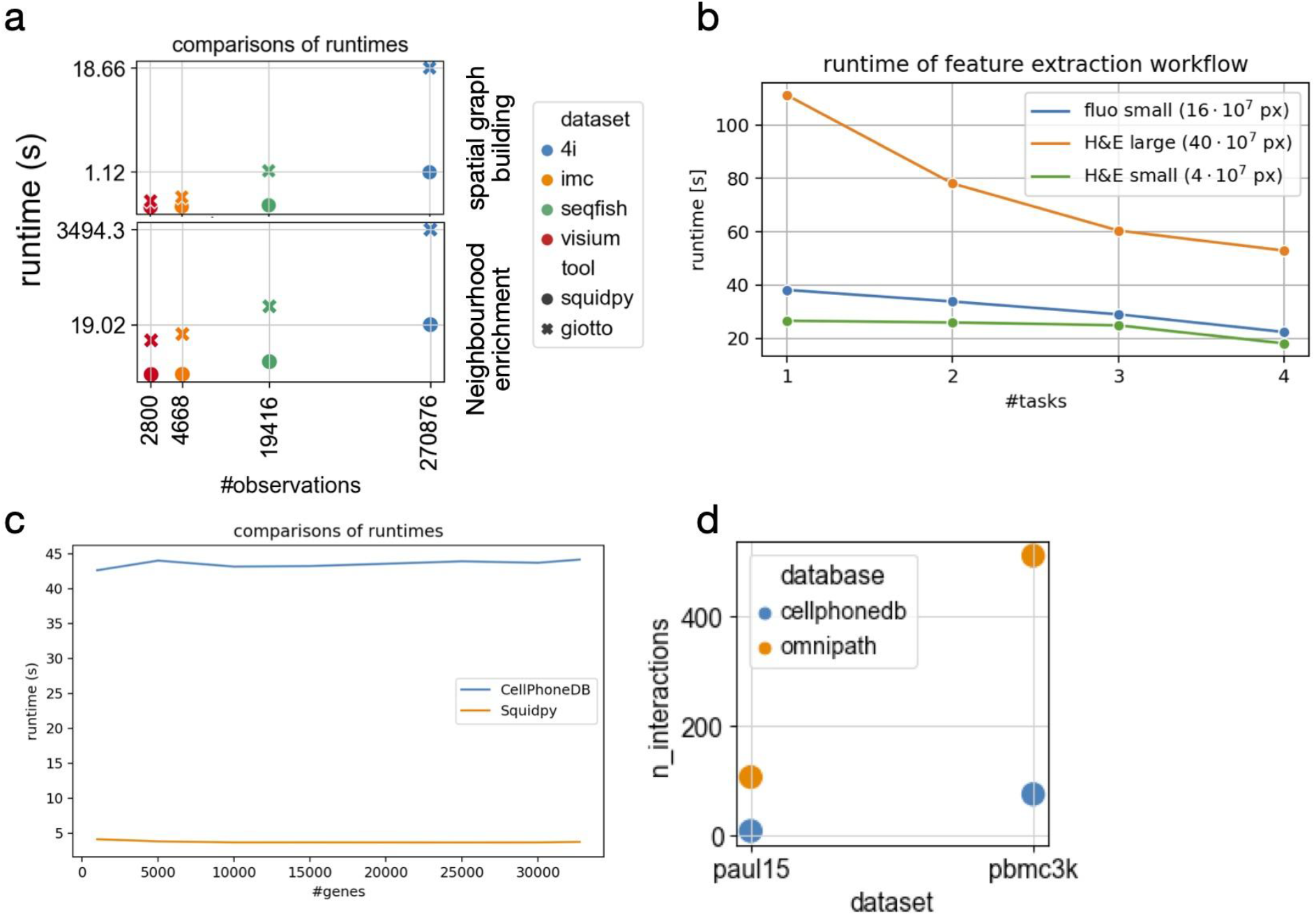
Benchmarking resources for Squidpy analysis modules. Benchmarks (a) and (b) were run on a 2,4 GHz Intel Core i5 processor with 4 cores and 16 GB RAM. Benchmarks (c) and (d) were run on a Centos 8 server cluster with 32 cores and 128 GB of memory. Unless explicitly mentioned, functions were run without parallelization. (a) Execution times for spatial graph building and neighborhood enrichment analysis, comparing four spatial datasets at increasing number of observations. Squidpy outperforms similar functions provided by the Giotto toolkit^11^, for any dataset and task. Reported are mean values for 10 runs, except for the 4i neighbor enrichment test that was run only once in Giotto. (b) Execution time for typical feature extraction workflow on different datasets. The feature extraction workflow consisted of segmenting the image using watershed with a fixed threshold, and extracting summary and segmentation features with default parameters. The segmentation was done using image tiles of size 2000. Using more cores (tasks) linearly decreases computation time for the feature extraction workflow, enabling processing of very large images (>400M pixels). (C) Execution time for Squidpy’s implementation of the CellphoneDB permutation-based test, at an increasing number of genes for the development of human forebrain dataset^37^. (d) Squidpy implementation of the CellphoneDB permutation-based tests uses the full Omnipath database for ligand receptor annotations. For two datasets (paul15^38^ mouse and pbmc3k^39^ human), Omnipath in Squidpy can recover a higher number of interactions.

**Supplementary Figure 2.**
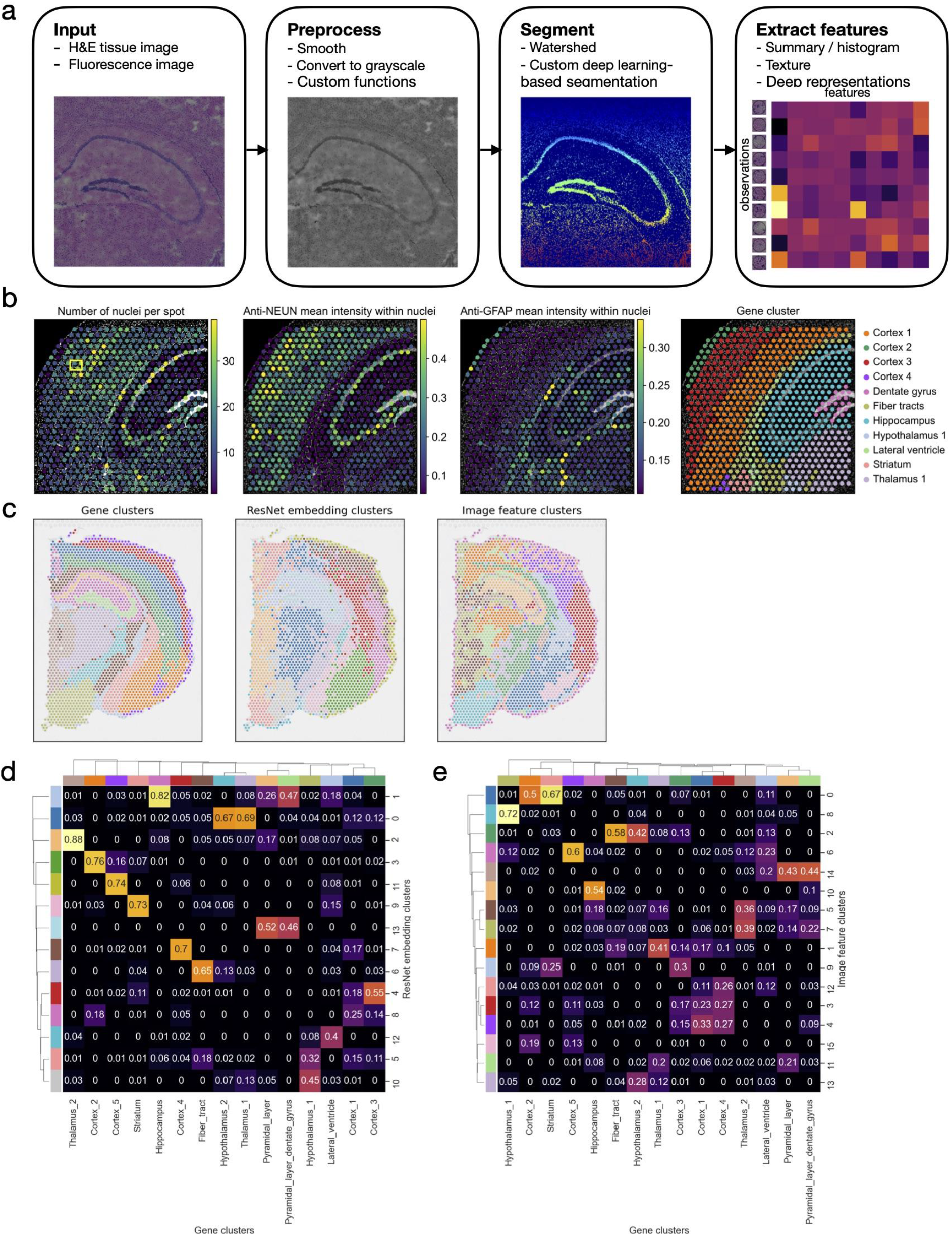
Image processing workflow and examples of segmentation and deep learning interface. (a) Exemplary image processing workflow utilising Squidpy’s Image Container object. From left to right are shown: the high-resolution source image, the preprocessing results (smooth, gray methods), the cell-segmentation results (that can be done with a watershed or custom, deep learning-based approach) and finally the feature extraction results. The features can be computed both at spot level, or at segmentation mask level, enabling the analysis to relate any pixel-level metric to the molecular profile. (b) Segmentation features extracted using a watershed segmentation. Extension of Figure 2 (a). From left to right are shown: number of nuclei underneath each Visium spot, mean intensity of anti-NEUN channel within the nuclei masks, mean intensity of anti-GFAP channel within the nuclei masks, and a leiden clustering of the gene expression values. The segmentation features provide interpretable, additional information to the gene-space clustering. We can see that the cell-rich pyramidal layer of the Hippocampus has more cells than the surrounding areas. This fine-grained differentiation of the Hippocampus is not visible in the gene clusters, where the Hippocampus is only one cluster. The per-channel intensities show that the areas labelled with “Cortex_1” and “Cortex_3” have a higher intensity of neurons (higher intensity of anti-NEUN channel) and that clusters “Fiber_tracts” and “lateral ventricles” are enriched with glial cells (higher intensity of anti-GFAP channel). (c) Qualitative comparison of gene-space clustering (left) with clustering of ResNet features (center) and clustering of summary features (right, see Fig. 2b) using a mouse brain Visium dataset with an H&E microscopy image. ResNet features were calculated by training a pre-trained ResNet model to predict the gene-expression cluster assignment (shown on the left) and taking the feature vector of the last fully connected layer as data representation. (d) Confusion matrix showing the proportion of assigned labels in gene clusters and resnet embedding clusters from (c). Rows correspond to clusters in gene expression space (c) left), columns correspond to resnet embedding clusters (c) center). The heatmap shows the proportion of overlapping observations in each cluster annotation. For instance, for “Thalamus_2” cluster, 88% of observations are annotated as cluster 2 in the resnet embedding visualization. We can see that for some cluster labels the prediction was strong, whereas for others the resnet model was unable to discriminate the labels. For instance, some regions of the cortex and hypothalamus seemed to not have been accurately classified. This showcases how the image container object can be used to relate morphology information from the source image to any annotation in the Anndata object. (e) Confusion matrix showing the proportion of assigned labels in gene clusters and image summary feature clusters from (c). Rows correspond to clusters in gene expression space (c) left), columns correspond to image summary feature clusters (c) center). The heatmap shows the proportion of overlapping observations in each cluster annotation. Several of the gene clusters are recognizable using simple image features. E.g.,“Hypothalamus_1” is overlapping to 77% with cluster 8, “Hippocampus” is overlapping to 54% with cluster 10, and “Pyramidial_layer” and“Pyramidial_layer_dentate_gyrus” are covered to 43%/44% by cluster 14. In other regions, especially the cortex (clusters “Cortex_1”, “Cortex_3”, “Cortex_4”), the image clusters do not overlap well (no cluster overlap > 33%), showing that in these regions simple image features and gene expression values show different patterns.

**Supplementary Figure 3.**
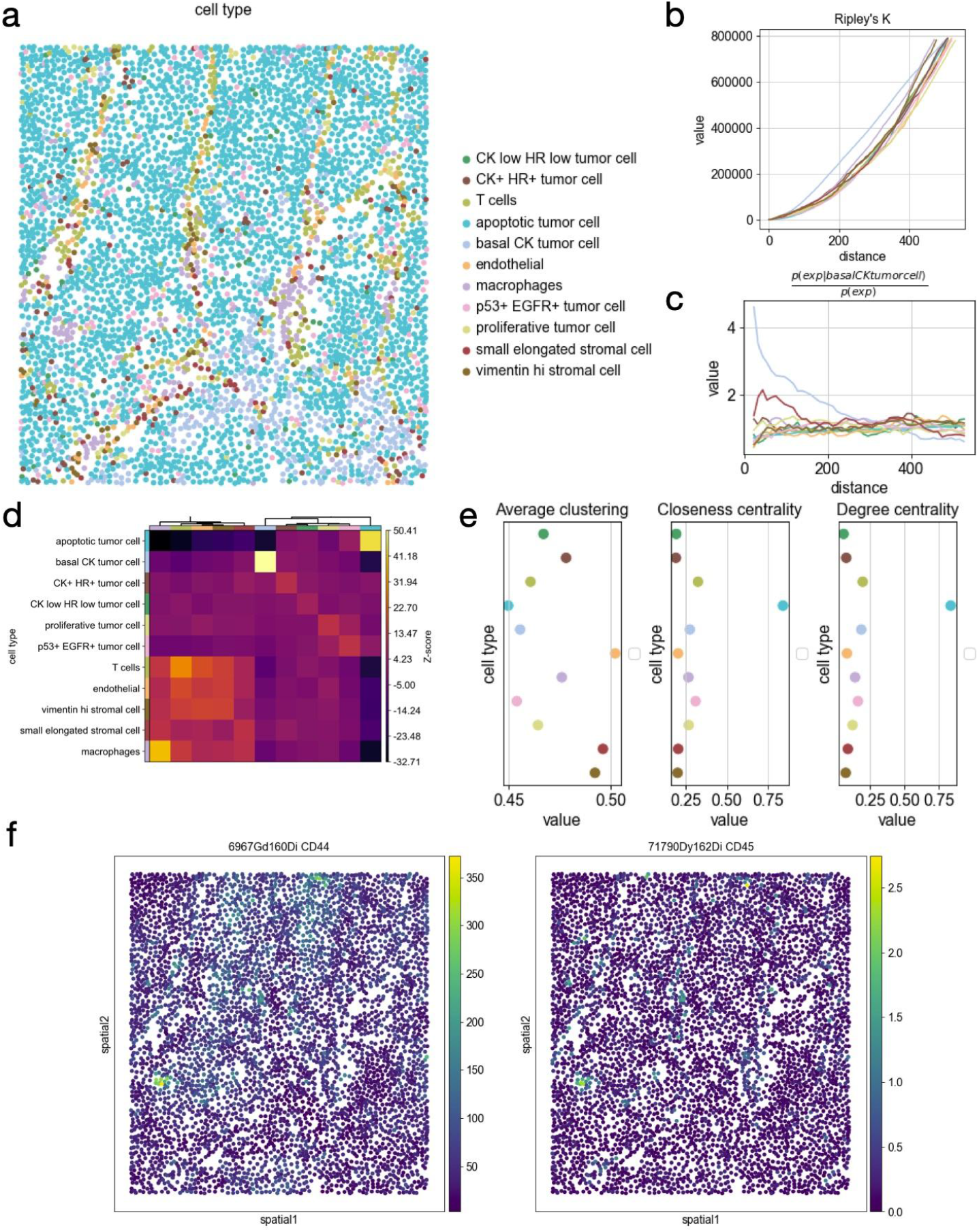
Example analysis of Imaging Mass Cytometry data from breast cancer biopsies. (a) Spatial visualization of cell types as defined by the original authors^33^. (b) Ripley’s K statistics computed at increasing distances threshold across the tissue. There is no clear spatial pattern in the data, except for a small increased clustering pattern of the “basal CK tumor cell”, which can be visualized in the lower-right section of the spatial plot. (c) Co-occurrence analysis of cell types at increasing distance thresholds across the tissue. Visualized is the probability conditioned on the presence of the “basal CK tumor cell”. Interestingly, we can observe a slight co-enrichment with the “small elongated stromal cell” cluster. (d) Neighborhood enrichment analysis between cell type clusters in the spatial graph. We can observe how the immune cell subsets and stromal cells seem to form a closer neighborhood as opposed to the tumor cells. (e) Network centralities for cell types (nodes of the spatial graph). The “apoptotic tumor cell” cluster shows high closeness and degree centrality, and it is indeed the most abundant and spread class label in the graph. (f) Visualization of two markers for immune cell populations, visualized in spatial coordinates.

**Supplementary Figure 4.**
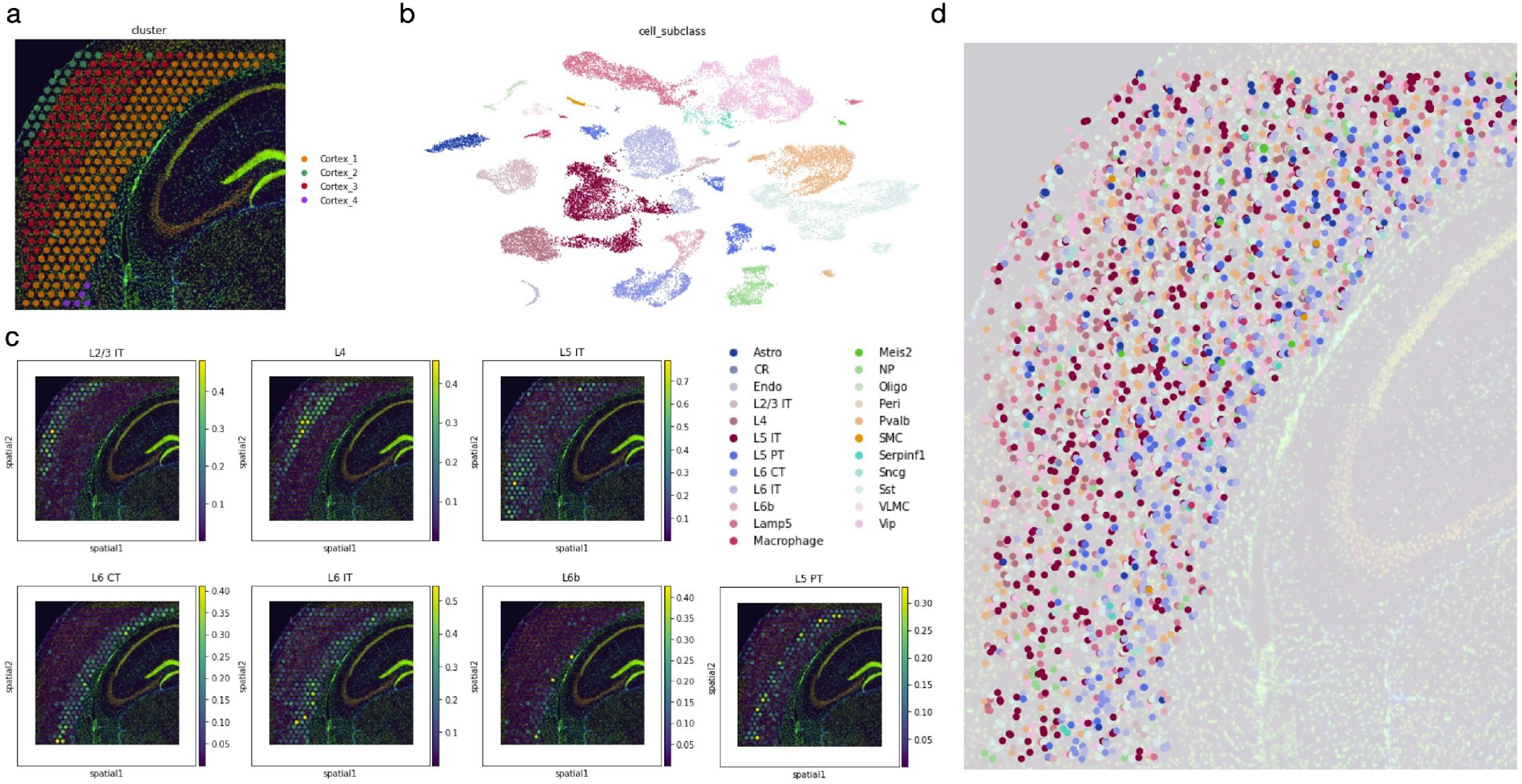
Interfacing Squidpy to Tangram for segmentation-aware cell-type deconvolution. Tangram is a recently published cell-type deconvolution method that maps single cell to spatial voxels of gene expression profiles. Squidpy’s Image Container can be used to acquire nuclei segmentation mask and leverage this mask to map cell types to tissue using Tangram. (a) Subset of Visium spatial transcriptomics dataset showing a mouse brain coronal section. (b) scRNA-seq data from the mouse cortex from Tasic et al^40^. (c) Tangram results as averaged by cell type. The cortical layers have been deconvoluted successfully. (d) Tangram maps of single cells. The cell type of the segmentation objects were assigned by Tangram, employing the seamless integration provided by Squidpy between the segmentation objects and the original spot observations in Anndata. In the figure, each point corresponds to a segmentation object colored by the cell type assigned by Tangram.

**Supplementary Table 1.**
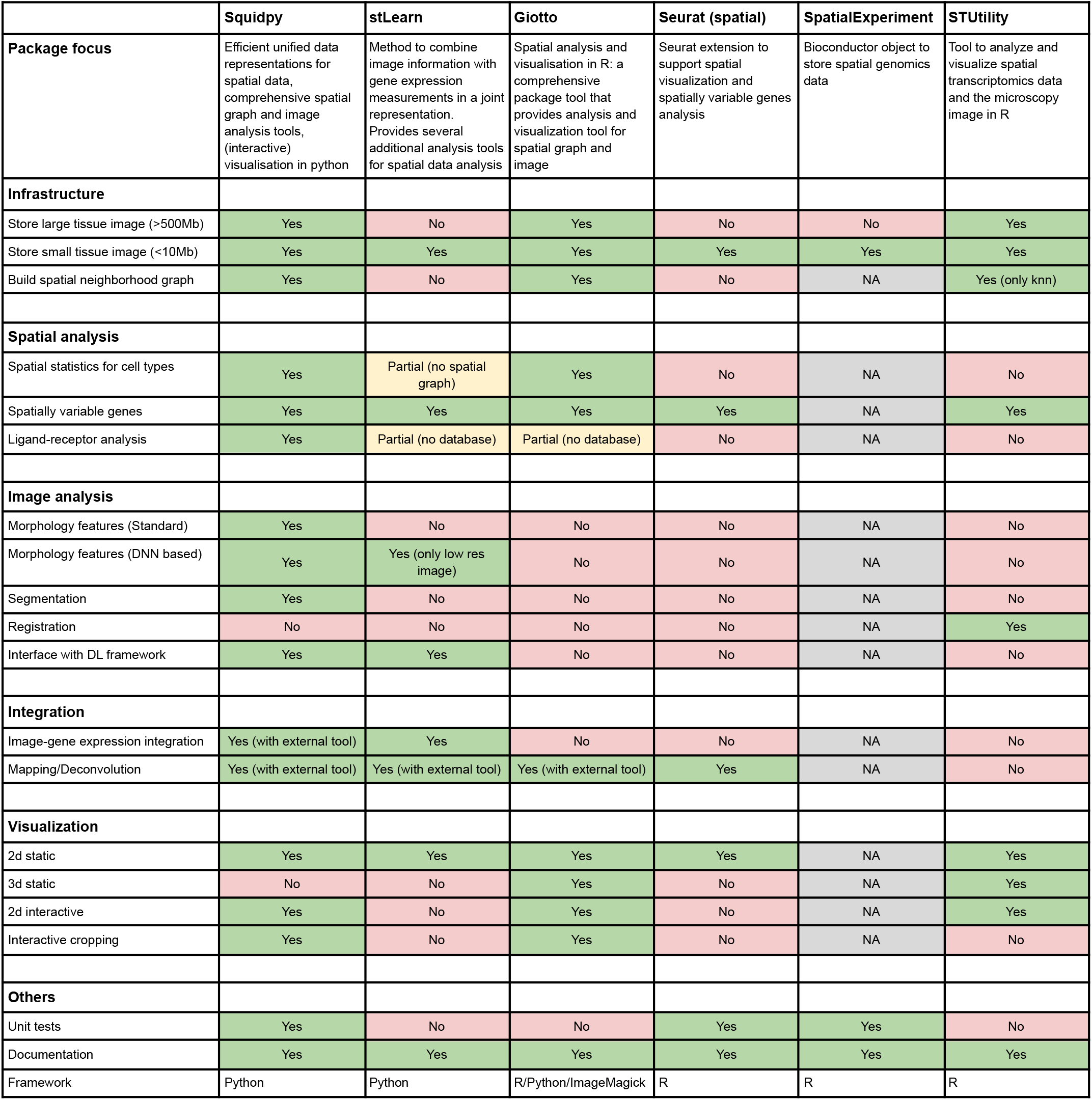
Comparison of Squidpy features to existing tools for spatial molecular data analysis. Rows correspond to a set of analysis features that are specific for working with spatial molecular data. It is subdivided in Infrastructure, Spatial Analysis, Image Analysis, Integration, Visualization and Others. The columns contain software tools that are tailored for spatial data analysis. Entries have been labelled according to whether the software tool is able to provide a specific functionality, whether it’s partially available or whether it’s missing. The row “Framework” specifies which programming languages are necessary to use all of the functionalities of the package. Finally, for SpatialExperiment, since it is an object to store spatial transcriptomics data, the analysis features do not apply.

## Squidpy - Online Methods

## Online methods

### 1. Infrastructure

#### Spatial graph

The spatial graph is a graph of spatial neighbors with cells (or spots in case of Visium) as nodes and neighborhood relations between spots as edges. We use spatial coordinates of spots to identify neighbors among them. Different approach of defining a neighborhood relation among spots are used for different types of spatial datasets.

Visium spatial datasets have a hexagonal outline for their spots, i.e each spot has up to eight spots situated around it. For this type of spatial dataset the parameter `n_rings` should be used. It specifies for each spot how many hexagonal rings of spots around it will be considered neighbors.

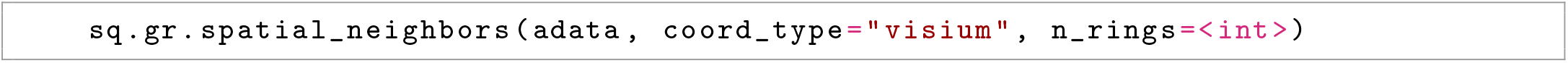

For the other types of spatial datasets neighbors can be defined as the closest spots in terms of euclidean distance between their coordinates. For a fixed number of the closest spots for each spot, it leverages the k-nearest neighbors search from Scikit-learn^1^ and `n_neigh` must be used.

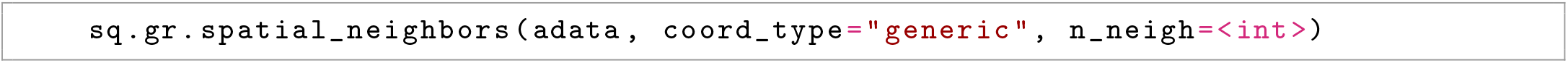

In order to get all spots within a specified radius (in units of the spatial coordinates) from each spot as neighbors, the parameter `radius` should be used.

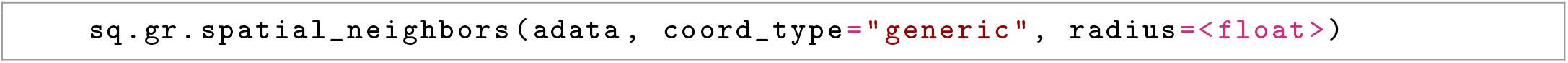

The function builds a spatial graph and saves its adjacency and weighted adjacency matrices to `adata.obsp[`’`spatial_connectivities`’] in either Numpy^2^ or Scipy sparse arrays ^3^. The weights of the weighted adjacency matrix are distances in the case of `coord_type=`“`generic`” and ordinal numbers of hexagonal rings in the case of `coord_type=`“`visium`”. Together with the connectivities, we also provide a sparse adjacency matrix of distances, saved in `adata.obsp[`’`spatial_distances` ’] We also provide spectral and cosine transformation of the adjacency matrix for uses in graph convolutional networks ^4^.

#### Image Container

The Image Container is an object for microscopy tissue images associated with spatial molecular datasets. The object is a thin wrapper of an `xarray.Dataset` ^5^ and provides efficient access to in-memory and on-disk images. On-disk files are loaded lazily using `dask` ^6^ through `rasterio` ^7^, meaning content is only read in memory when requested. The object can be saved as a zarr store `zarr` ^8^. This allows handling very large files that do not fit in memory.

Image Container is initialised with an in-memory array or a path to an image file on disk. Images are saved with the key `layer`. If lazy loading is desired, the `chunks` parameter needs to be specified.

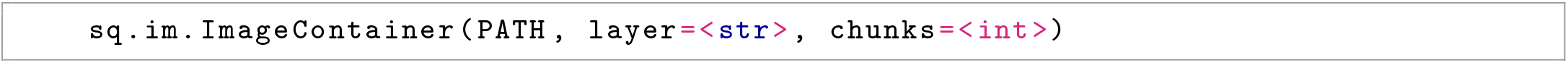

More images layers with the same spatial dimensions `x` and `y` like segmentation masks can be added to an existing Image Container.

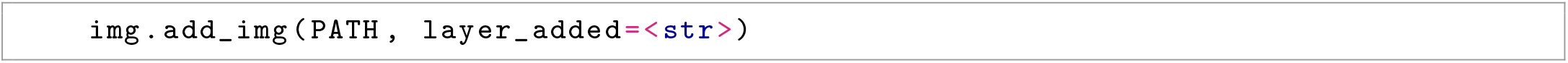

The Image Container is able to interface with Anndata objects, in order to relate any pixel-level information to the observations stored in Anndata (e.g. cells, spots etc.). For instance, it is possible to create a generator that yields image’s crops on-the-fly corresponding to locations of the spots in the image:

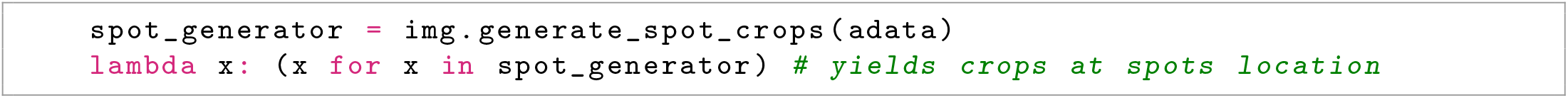

This of course works for both features computed at crop-level but also at segmentation-object level. For instance, it is possible to get centroids coordinates as well as several features of the segmentation object that overlap with the spot capture area.

#### Napari for interactive visualization

Napari is a fast, interactive, multi-dimensional image viewer in Python^9^. In squidpy, it is possible to visualize the source image together with any anndata annotation with Napari. Such functionality is useful for fast and interactive exploration of analysis results saved in anndata together with the high resolution image. Furthermore, leveraging Napari functionalities, it is possible to manually annotate tissue areas and assign underlying spots to annotations saved in the Anndata object. Such ability to relate manually defined tissue areas to observations in anndata is particularly useful in settings where there is a pathologist annotation available and it needs to be integrated with analysis at gene expression or image level. All the steps described here are done in Python, therefore available in the same environment where the analysis is performed (it does not require an additional tool).

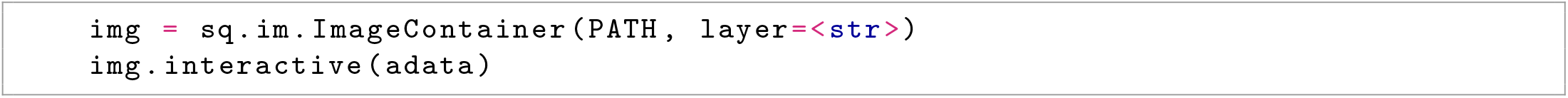

### 2. Graph and spatial patterns analysis

#### Neighborhood enrichment test

The association between label pairs in the connectivity graph is estimated by counting the sum of nodes that belong to classes i and j (e.g. cluster annotation) and are proximal to each other, noted *x*_*ij*_. To estimate the deviation of this number versus a random configuration of cluster labels in the same connectivity graph, we scramble the cluster labels while maintaining the connectivities, and then recount the number of nodes recovered in each iteration (1,000 times by default). Using these estimates, we calculate expected means (*µ*_*ij*_) and standard deviations (*σ*_*ij*_) for each pair, and a Z-score as,

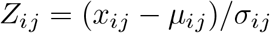

The Z-score indicates if a cluster pair is over-represented or over-depleted for node-node interactions in the connectivity graph. This approach was first described (to the best of our knowledge) by Schapiro et al ^10^. The analysis and visualization can be performed with the analysis code showed below.

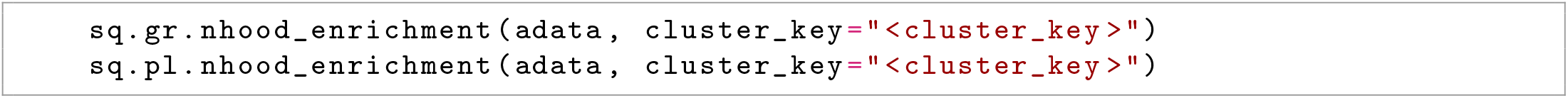

Our implementation leverages just-in-time compilation with Numba ^11^ to achieve greater performances in computation time (see **Supplementary figure 1**).

#### Ligand-receptor interaction analysis

We provide a re-implementation of the popular Cell-phonedb method for ligand-receptor interaction analysis ^12^. In short, it’s a permutation-based test of ligand-receptor expression across cell-types combinations. Given a list of annotated ligand-receptor pairs, the test computes the mean expression of the two molecules (ligand, receptor) between cell types, and builds a null-distribution based on *n* permutations (default 1000). A p-value is computed based on the proportion of the permuted means against the true mean. In Cellphonedb, if a receptor or ligand is composed of several subunits, the minimum expression is considered for the test. In our implementation, we also include the option of taking the mean expression of all molecules in the complex. Our implementation also employs Omnipath ^13^ as ligand-receptor interaction annotatiojn. A larger database that contains the original CellphoneDB database together with 5 other resources (see Turei et al. ^13^). Finally, our implementation leverages just-in-time compilation with Numba ^11^ to achieve greater performances in computation time (see **Supplementary figure 1**).

#### Ripley’s K function

is a spatial analysis method used to describe whether points with discrete annotation in space follow random, dispersed or clustered patterns. Ripley’K function can be used to describe the spatial patterning of cell clusters in the area of interest. Ripley’s K function is defined as

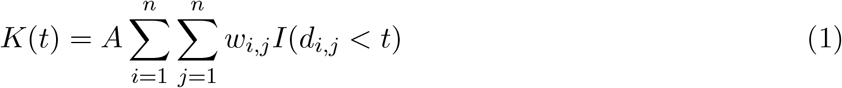

Where *I*(*d*_*i,j*_ *< t*) is the indicator function, that sets whether the operand is 1 or 0 based on the (euclidean) distance *d*_*i,j*_ evaluated at search radius *t, A* is the average density of point in the area of interest and *w*_*i,j*_ is the edge effect correction (see Astropy implementation for details on this term^14^).

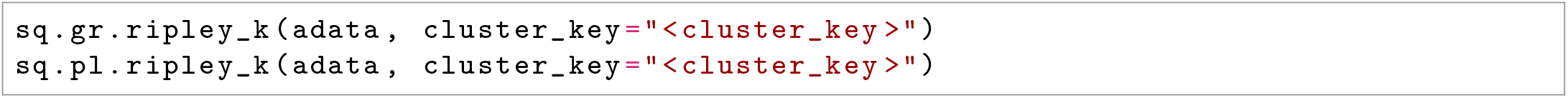

#### Cluster co-occurrence ratio

provides a score on the co-occurrence of clusters of interest across spatial dimensions. It is defined as

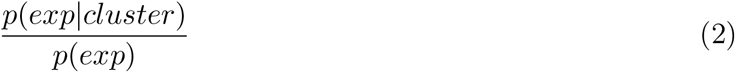

where *cluster* is the annotation of interest to be used as conditioning for the co-occurrence of all clusters. It is computed across *n* radius of size *d* across the tissue area. It was inspired by an analysis performed by Tosti et al. to investigate tissue organization in the human pancreas with spatial transcriptomics^15^.

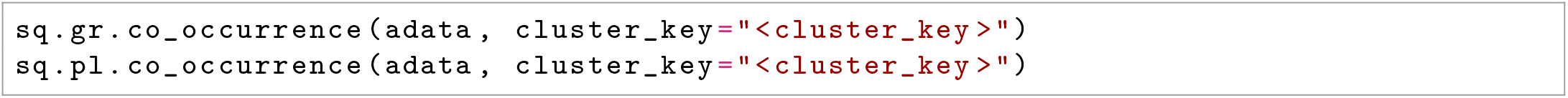

#### Global Moran’s I

is a spatial auto-correlation statistics, widely used in spatial data analysis. Given a feature (gene) and spatial location of observations, it evaluates whether the pattern expressed is clustered, dispersed, or random^16^. It is defined as:

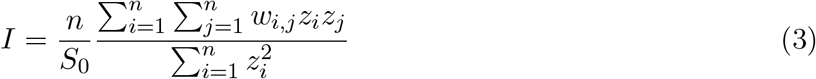

where *z*_*i*_ is the deviation of the feature from the mean 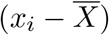, *w*_*i,j*_ is the spatial weight between observations, *n* is the number of spatial units. We provide an wrapper for the global Moran’s I

statistics implemented in libpysal ^17^. Test statistics and p values (computed from a permutation based test and further FDR corrected) are stored in `adata.uns[`“`moranI`”].

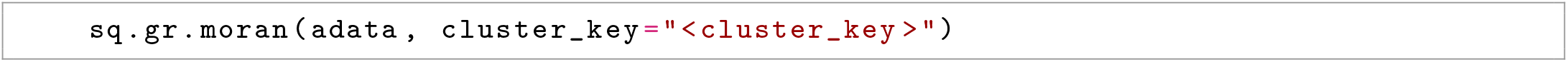

#### Centrality scores

provide a numerical analysis on node patterns in the graph, which helps to better understand complex dependencies in large graphs. A centrality is a function *C* which assigns every vertex *v* in the graph a numeric value *C*(*v*) ∈ ∝. It therefore gives a ranking of the single components (i.e. cells) in the graph which simplifies to identify key individuals. Group centrality measures have been introduced by Everett and Borgatti. ^18^. They provide a framework to assess clusters of cells in the graph, i.e. is a specific cell type more central or more connected in the graph than others. Let *G* = (*V, E*) be a graph with nodes *V* and edges *E*. Additionally, let *S* be a group of nodes allocated to the same cluster *c*_*S*_. Then *N* (*S*) defines the neighbourhood of all nodes in *S*. The following four (group) centrality measures are implemented. **Group degree centrality** is defined by the fraction of non-cluster members that are connected to cluster members, so

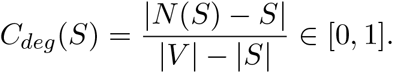

Larger values indicate a more central cluster. Group degree centrality can help to identify essential clusters or cell types in the graph. **Group closeness centrality** measures how close the cluster is to other nodes in the graph and is calculated by the number of non-group members divided by the sum of all distances from the cluster to all vertices outside the cluster, so

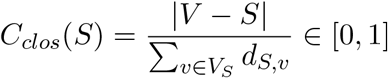

where *d*_*S,v*_ = min_*u∈S*_ *d*_*u,v*_ is the minimal distance of the group *S* from *v*. Hence, larger values indicate a greater centrality. **Group betweenness centrality** measures the proportion of shortest paths connecting pairs of non-group members that pass through the group. Let *S* be a subset of a graph with vertex set *V*_*S*_. Let *g*_*u,v*_ be the number of shortest paths connecting *u* to *v* and *g*_*u,v*_(*S*) be the number of shortest paths connecting *u* to *v* passing through *S*. The group betweeenness centrality is then given by

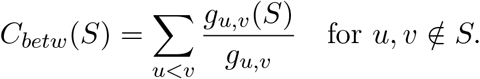

The properties of this centrality score are fundamentally different from degree and closeness centrality scores, hence results often differ. The last measure described is the **average clustering coefficient**. It describes how well nodes in a graph tend to cluster together. Let *n* be the number of nodes in *S*. Then the average clustering coefficient is given by

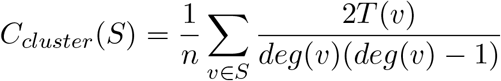

with *T* (*v*) being the number of triangles through node *v* and *deg*(*v*) the degree of node *v*. The describes centrality scores have been implemented using the NetworkX library in python^19^.

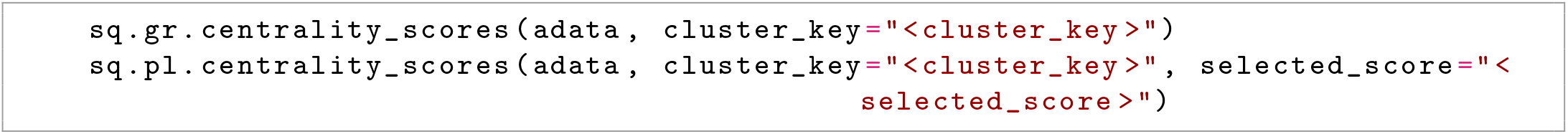

#### Interaction matrix

represents the total number of edges that are shared between nodes with specific attributes (e.g. clusters or cell types).

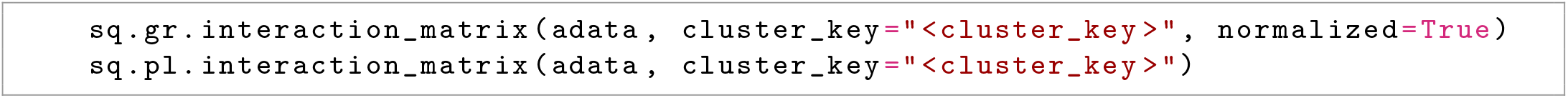

Python implementations relies ont the NetworkX library^19^.

### 3. Image analysis and segmentation

#### Image processing

Before extracting features from microscopy images, the images can be preprocessed. Squidpy implements functions for commonly used preprocessing functions like conversion to gray-scale or smoothing using a gaussian kernel.

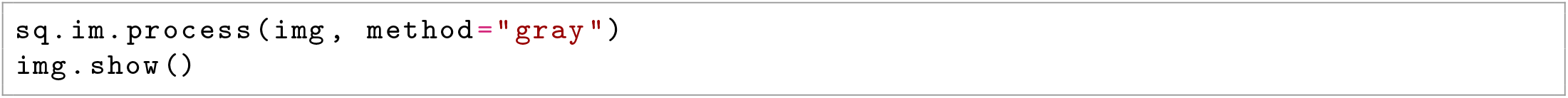

Implementations are based on the Scikit-image package ^20^ and allow processing of very large images through tiling the image into smaller crops and processing these.

#### Image segmentation

Nuclei segmentation is an important step when analysing microscopy images. It allows the quantitative analysis of the number of nuclei, their areas, and morphological features. There are a wide range of approaches for nuclei segmentation, from established techniques like thresholding to modern deep learning-based approaches.

A difficulty for nuclei segmentation is to distinguish between partially overlapping nuclei. Watershed is a classic algorithm used to separate overlapping objects by treating pixel values as local topology. For this, starting from points of lowest intensity, the image is flooded until basins from different starting points meet at the watershed ridge lines.

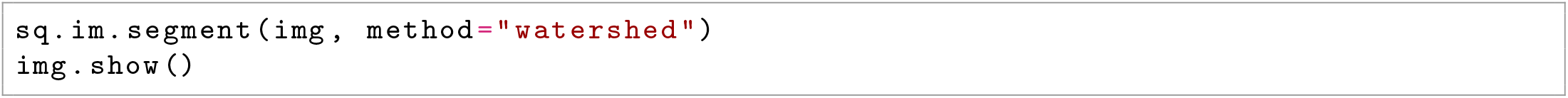

Implementations in Squidpy are based on the original Scikit-image python implementation^20^.

#### Custom approaches with deep learning

Depending on the quality of the data, simple segmentation approaches like watershed might not be appropriate. Nowadays, many complex segmentation algorithms are provided as pre-trained deep learning models, such as Stardist^21^, Splinedist^22^ and Cellpose ^23^. These models can be easily used within the segmentation function.

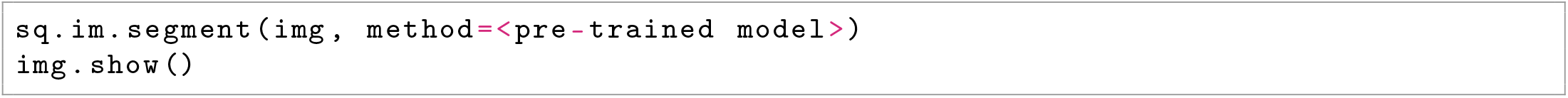

#### Image features

Tissue organisation in microscopic images can be analysed with different image features. This filters relevant information from the (high dimensional) images, allowing for easy interpretation and comparison with other features obtained at the same spatial location. Image features are calculated from the tissue image at each location (*x, y*) where there is transcriptomics information available, resulting in a obs x features features matrix similar to the obs x gene matrix. This image feature matrix can then be used in any single-cell analysis workflow, just like the gene matrix.

The scale and size of the image used to calculate features can be adjusted using the `scale` and `spot_scale` parameters. Feature extraction can be parallelized by providing `n_jobs` (see Supplementary Figure 1). The calculated feature matrix is stored in `adata[key]`.

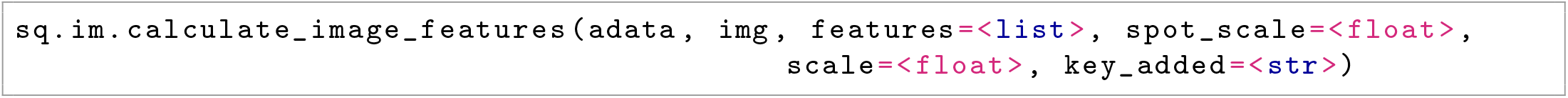

*Summary features* calculate the mean, the standard variation or specific quantiles for a color channel. Similarly, *histogram features* scan the histogram of a color channel to calculate quantiles according a defined number of bins.

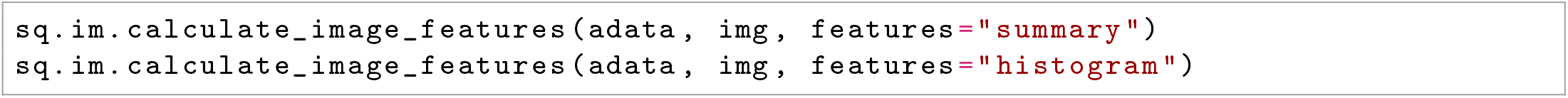

*Textural features* calculate statistics over a histogram that describes the signatures of textures. To grasp the concept of texture intuitively the inextricable relationship between texture and tone is considered^24^ : if a small-area patch of an image has little variation in it’s gray tone the dominant property of that area is tone. If the patch has a wide variation of gray tone features, the dominant property of the area is texture. An image has a simple texture if it consists of recurring textural features. For a grey level image **I** or e.g. a fluorescence color channel, a co-occurrence matrix **C** is computed. **C** is a histogram over pairs of pixels (*i, j*) with specific values (*p, q*) ϵ [0, 1, …, 255]^2^ and a fixed pixel offset:

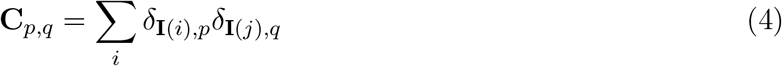

with Kronecker-Delta *δ*. The offset is a fixed pixel distance from *i* to *j* under a fixed direction angle. Based on the co-occurence matrix different meaningful statistics (*texture properties*) can be calculated which summarize textural pattern characteristics of the image:

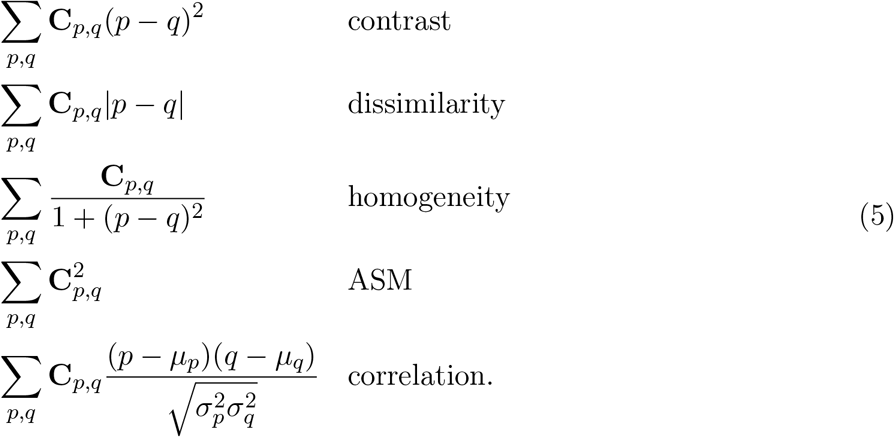

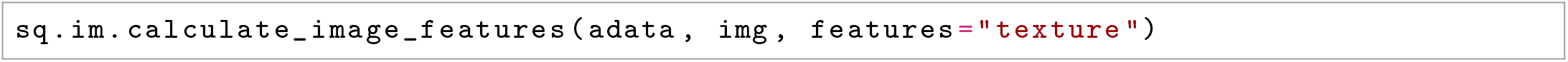

All the above implementations rely on the Scikit-image python package ^20^.

#### Segmentation features

Similar to *image features* that are extracted from raw tissue images, *segmentation features* can be extracted from a segmentation object (3). These features allow to get statistics over the number, area, and morphology of the nuclei in one image. To compute these features, the Image Container `img` needs to contain a segmented image at layer `<segmented_img>`

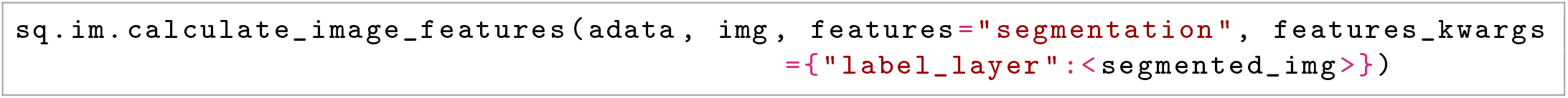

#### Custom features based on deep learning models

Squidpy feature calculation function can also be used with custom user-defined features extraction functions. This enables the use of e.g., pre-trained deep learning models as feature extractors.

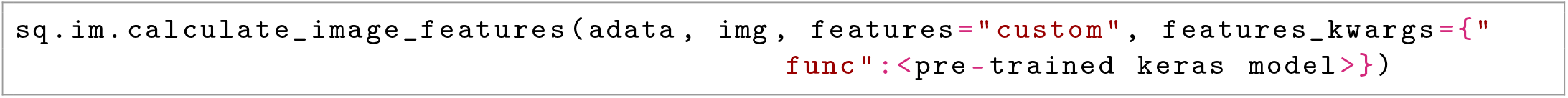

